# De novo assembly of *trachidermus fasciatus* genome by nanopore sequencing

**DOI:** 10.1101/2020.04.18.042093

**Authors:** Gangcai Xie, Xu Zhang, Feng Lv, Mengmeng Sang, Hairong Hu, Jinqiu Wang, Dong Liu

**Author notes:** Those authors contribute equally to this project (Gangcai Xie, Xu Zhang). Authors for correspondence: Gangcai Xie, Ph.D, Institute of Reproductive Medicine, Medical School, Nantong University, Qixiu Road 19, Nantong, China, 226001, Or Jinqiu Wang, Ph.D, School of Life Sciences, Institute of Genetics, State Key Laboratory of Genetic Engineering, Fudan University, Shanghai, China, 200433, Or Dong Liu, Ph.D, Jiangsu Key Laboratory of Neuroregeneration, Nantong University, Qixiu Road 19, Nantong, China, 226001.

## Abstract

*Trachidermus fasciatus* is a roughskin sculpin fish widely located at the coastal areas of East Asia. Due to the environmental destruction and overfishing, the populations of this species have been under threat. It is important to have a reference genome to study the population genetics, domestic farming, and genetic resource protection. However, currently, there is no reference genome for *Trachidermus fasciatus*, which has greatly hurdled the studies on this species. In this study, we proposed to integrate nanopore long reads sequencing, Illumina short reads sequencing and Hi-C methods to thoroughly de novo assemble the genome of *Trachidermus fasciatus*. Our results provided a chromosome-level high quality genome assembly with a total length of about 543 Mb, and with N50 of 23 Mb. Based on de novo gene prediction and RNA sequencing information, a total of 38728 genes were found, including 23191 protein coding genes, 2149 small RNAs, 5572 rRNAs, and 7816 tRNAs. Besides, about 23% of the genome area is covered by the repetitive elements. Furthermore, The BUSCO evaluation of the completeness of the assembled genome is more than 96%, and the single base accuracy is 99.997%. Our study provided the first whole genome reference for the species of *Trachidermus fasciatus*, which might greatly facilitate the future studies on this species.

## Introduction

Roughskin sculpin (*Trachidermus fasciatus Heckel*) is a small, carnivorous and catadromous fish that had been found in China, Korean and Japan coastal areas[1, 2]. Historically, roughskin sculpin had been named as one of the four most famous fishes in China, and had been treated as valuable food sources by Chinese[3]. However, due to overfishing, and the environmental changes of spawning and habitat sites, population size of roughskin sculpin had declined significantly during past decades[1, 4]. Since 1988, roughskin sculpin had been listed as Class II protected animal by Chinese government, which encouraged our development of the farming system for roughskin sculpin domestication[5]. Recently, genetic diversity[6] and genomic signature[7] had been studied for *Trachidermus fasciatus*, which might be important for its conservation management. For both of farming system development and genetic studies of roughskin sculpin, it would be important to have a reference genome of this species. Although the mitochondrial genome had been completed previously[8], the whole genome sequence information is still unavailable.

During past decades, high throughput DNA sequencing technologies had been advanced significantly, including Illumina short reads sequencing, Pacific biosciences and Oxford nanopore long reads sequencing[9, 10]. It had been illustrated that nanopore long reads sequencing technology can be used to assemble genomes from different species, including bacterial[11], human[12, 13], and rice[14]. It had been proved that chromosome-scale assemblies of human and mouse genomes can be generated by integrating short-reads DNA sequencing and Hi-C chromatin interaction mate-pair sequencing[15]. Through combining Hi-C and short reads data, 99% scaffolds spatial orienting accuracy was achieved[15].

In this study, Oxford nanopore sequencing, Illumina short reads sequencing technologies and the Hi-C method were integrated for de novo assembling *Trachidermus fasciatus* genome. This study provided the first high quality reference genome for the communities studying roughskin sculpin, and such information might greatly facilitate future studies on this species, including gene editing based selective breeding.

## Results

### Sequencing data sets

In order to get a high-quality genome assembly for *Trachidermus fasciatus*, we integrated long reads nanopore sequencing, high-quality short reads Illumina sequencing technologies and Hi-C method (Figure 1). For nanopore sequencing (Table 1), we got about 4 million reads passing quality control, containing more than 87 billion nucleotide bases. The length of longest reads is more than 240kb, and the N50 of all the reads is about 30kb.More than 70% of the reads with a length larger than 10kb, and about 12% of the reads with a length larger than 40kb.For Illumina sequencing, we got more than 350 million reads for both of genome and Hi-C sequencing (Table 1),which containing more than 50 billion nucleotide bases respectively. On average, nanopore reads is more than 140 times longer than Illumina reads.

**Figure 1:**
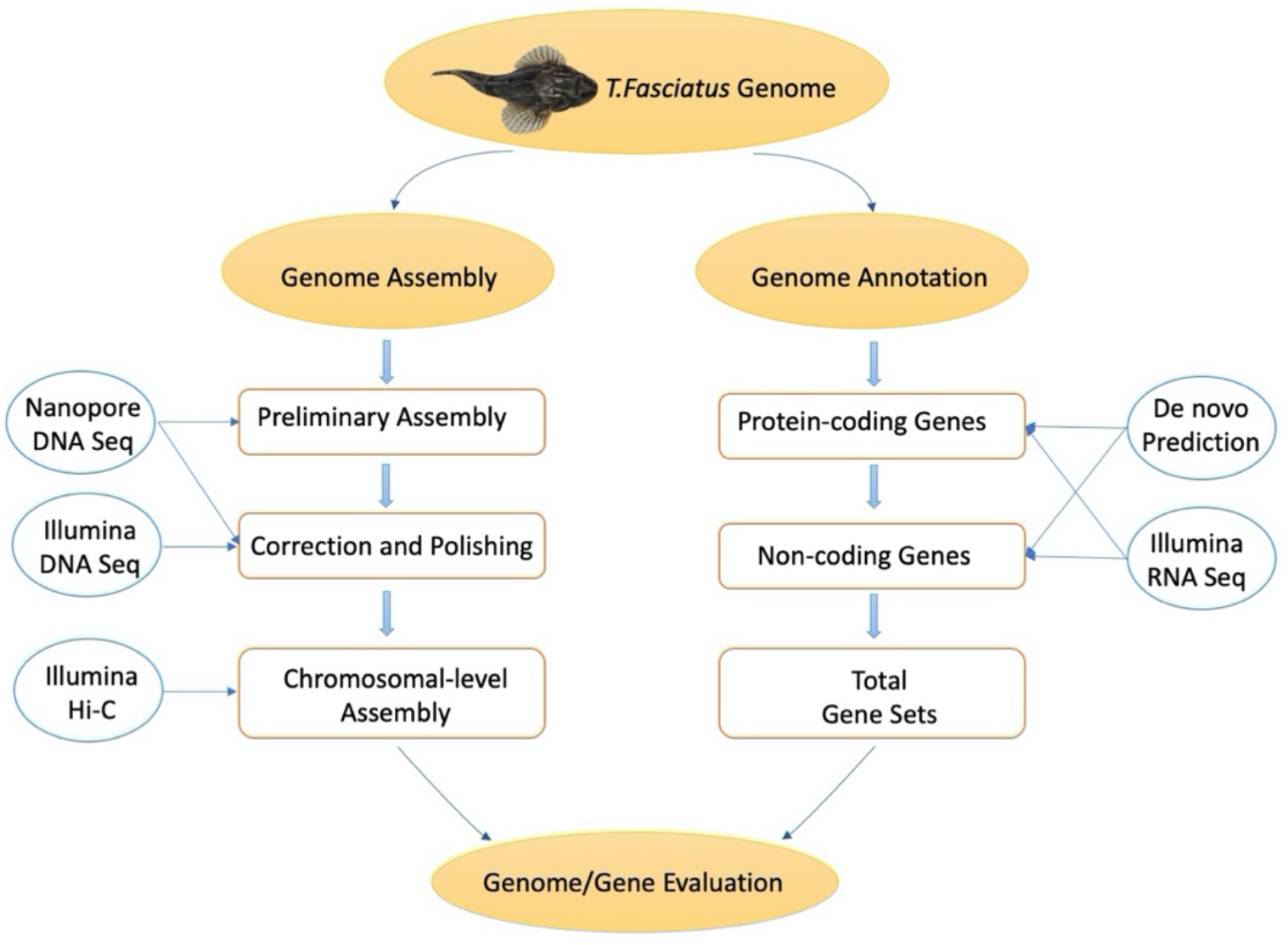
Pipeline of *T.Fasciatus* Genome assembly and annotation.

**Table 1:** Summary of sequencing data sets.

| library | # of bases | # of reads | reads length (mean, bp) | reads length(max, bp) |
| --- | --- | --- | --- | --- |
| nanopore-seq-lib1 | 45,424,703,117 | 2,049,727 | 22,161 | 240,976 |
| nanopore-seq-lib2 | 41,854,167,498 | 2,031,772 | 20,599 | 243,222 |
| Total | 87,278,870,615 | 4,081,499 | 21,384 | 243,222 |
| library | N50 | > 10kb percentage | > 20kb percentage | > 40kb percentage |
| nanopore-seq-lib1 | 31,576 | 75.85 | 48.11 | 13.48 |
| nanopore-seq-lib2 | 30,199 | 70.82 | 42.38 | 11.99 |
| Total | 30,943 | 73.35 | 45.26 | 12.74 |
| library | # of bases | # of reads | reads length (mean, bp) | reads length(max, bp) |
| NGS-genome | 54,380,742,900 | 362,538,286 | 150 | 150 |
| NGS-HiC | 60,291,587,700 | 401,943,918 | 150 | 150 |

### De novo assembly and genome polishing

*Trachidermus fasciatus* genome was preliminarily assembled based on nanopore sequencing data, and then polished based on both of nanopore and Illumina sequencing data. At the preliminary stage,62 contigs were assembled, the N50 is more than 23 Mbp (Million base pairs), and the longest contigs for the preliminary assembly is more than 35 million bp, while the total length of the preliminary genome is 539,115,043 bp (Table 2). After polishing, the N50 of the assembly increases from 23.4 Mbp to 23.55 Mbp, and the total size of polished assembly is about 542.6 Mbp. Based on N90 information (Table 2), 23 contigs can have a combined length covering 90% of the genome area, and this number is close to the number of chromosome pairs reported previously (karyotype 2N=40 by the studies of Jianhua *et al.[16]* and Jinqiu Wang[3]).

**Table 2:** Summary about the primary genome assembly

|  | Assembly (Preliminary) |  | Assembly (Polished) |  |
| --- | --- | --- | --- | --- |
|  | Contig Length(bp) | Contig Number | Contig Length(bp) | Contig Number |
| N50 | 23,408,022 | 10 | 23,556,738 | 10 |
| N60 | 20,522,422 | 13 | 20,673,841 | 13 |
| N70 | 18,786,353 | 16 | 18,887,485 | 16 |
| N80 | 17,637,255 | 18 | 17,756,054 | 18 |
| N90 | 7,606,484 | 23 | 7,646,988 | 23 |
| Longest | 35,774,240 | 1 | 36,041,605 | 1 |
| Total | 539,115,043 | 62 | 542,654,729 | 62 |
| Length>=1kb | 539,115,043 | 62 | 542,654,729 | 62 |
| Length>=2kb | 539,115,043 | 62 | 542,654,729 | 62 |
| Length>=5kb | 539,115,043 | 62 | 542,654,729 | 62 |

### Genome quality evaluation

The quality of the assembled *trachidermus fasciatus* genome was further evaluated by different methods. Firstly, the GC content and nanopore sequencing depth distribution were examined based on 10kb sliding windows. As shown in Figure 2A, only one peak observed for the GC content distribution as well as sequencing depth distribution, which indicates no contamination from other species. Then, the completeness of the assembled genome was evaluated by both of CEGMA and BUSCO(Figure 2B,2C), which illustrated a high percentage of completeness (98.39% and 96.95% respectively).Thirdly, high quality Illumina sequencing reads were mapped onto the assembled genome to evaluate its quality, and the results showed that 99.48% of the reads can be mapped, and 99.35% of the assembled genome is covered by at least one time. Finally, Genome contamination was further examined by matching the assembled contigs with known metazoa genome sequences (50kb bins). As shown in Figure 2E, over 98% of the genome length can be matched with the known metazoan genome sequences, which indicates no significant contamination from bacteria.

**Figure 2:**
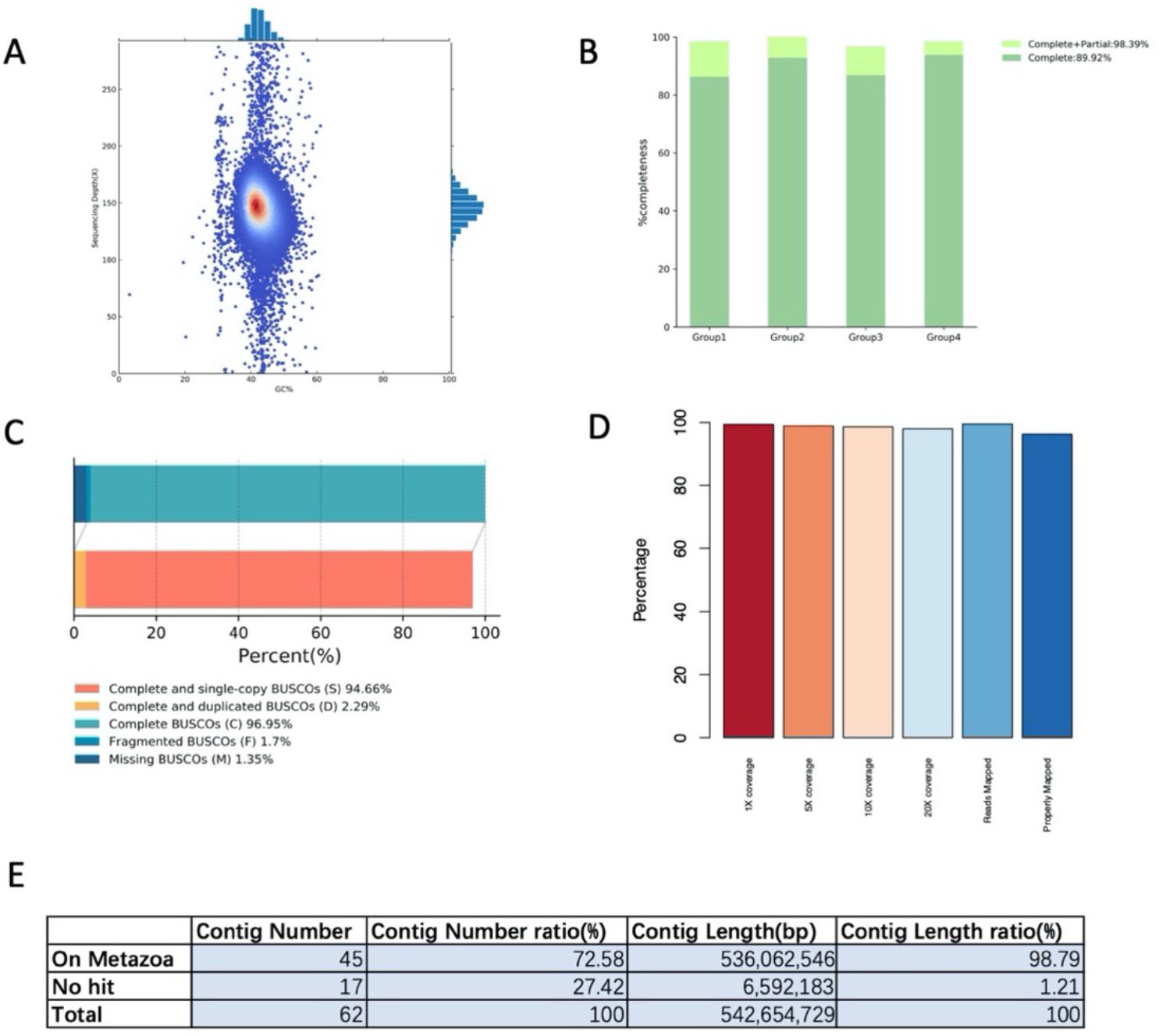
Genome assembly quality evaluation. (A)GC content and sequencing depth distribution. (B)CEGMA evaluation. (C) BUSCO evaluation. 94.66% coverage by completed and single-copy BUSCO sequences. (D) Illumina reads mappability and genome coverage. 99.35% of the assembled genome is covered by at least one time by illumina reads, 97.93% is covered by at least 20 times. And 99.48% of illumnina reads were mapped to the assembled genome, and 96.22% was properly mapped. (E) Genome contamination evaluation.

By mapping the Illumina short reads onto the assembly, the single base level accuracy was examined. At the depth of 5X genome-level coverage, 3444,14558 homogenenous SNPs and Indels were found respectively, which in total occupy 0.002683% of the genome. This information illustrated 99.996683% single base accuracy for the final assembled genome.

### Hi-C proximity-guided improvement of genome assembly

In the primary assembly based on DNA sequencing datasets, the number of disconnected contigs is 62. Comparing to the primary assembly, the Hi-C enhanced assembly showed cumulative longer scaffolds (Figure 3A). Using Hi-C chromatin interaction data, the contigs were re-arranged based on the interaction information, and in total 44 scaffolds were assembled (Figure 3B, C). Based on the top 20 longest scaffolds, which represent the main chromosomal scaffolds of *Trachidermus fasciatus*, the Hi-C chromatin interaction events were significantly enriched in the regions from the same chromosomes (Figure 3D). The number of N90 contigs reduced from 23 to 18 (primary assembly), and the top 20 scaffolds have a cumulative length of 533,710,745 bp, occupying 98.35% of the whole genome. This information indicates that the chromosome level genome assembly was completed after Hi-C proximity-guided improvement (Figure 3E).

**Figure 3:**
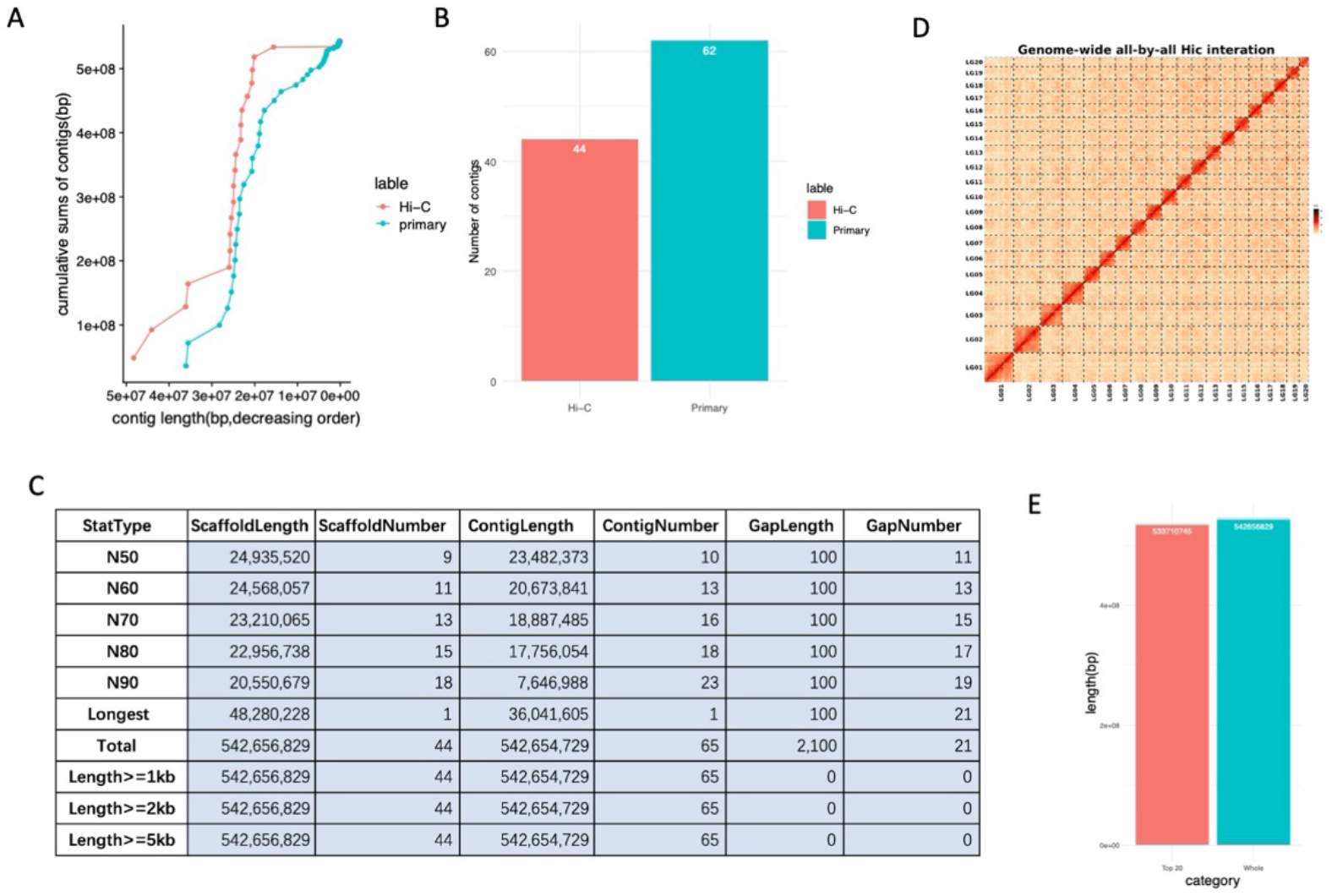
Hi-C enhanced genome assembly. (A)Comparison of cumulative sums of the contigs between primarily assembled genome and the Hi-C enhanced one. (B)Reduced number of scaffolds/contigs in Hi-C enhanced genome. (C)Statistics of the Hi-C enhanced assembly. (D)Genome-wide all-by-all Hi-C interaction heatmap based on top 20 scaffolds (main chromosomal scaffolds). (E) The cumulative length of top 20 longest scaffolds.

### *Trachidermus fasciatus* Genome Annotation

We integrated both of de novo and RNA sequencing based annotation methods for *Trachidermus fasciatus* genome annotation. Based on AUGUSTUS de novo protein coding gene prediction, 25,741 genes were found, based on GeMoMa homolog gene prediction, 22,211 genes were found, and based on RNA-seq data, 14,238 genes were discovered. Through the integration of the genes from those three methods by Evidence Modeler (EVM), 23,191 protein coding genes were finally identified for *Trachidermus fasciatus*. Besides protein coding genes, 5572,2149 and 7816 non-protein coding genes of rRNA,small RNA and tRNA respectively were also identified(Figure 4A).Comparing to the five close species of *Trachidermus fasciatus* with genome annotated, no abnormal length distribution of CDS, gene, exon and intron is observed(Figure 4B).Furthermore, the repetitive elements were also annotated, in total 23.7% of the *Trachidermus fasciatus* genome is covered by repetitive elements, including 4% LTR, 6.4% LINE,0.6% SINE and 7% DNA repeats(Figure 4C).There are about 157 thousands LINE elements, 526 thousands DNA elements, 166 thousands LTR elements and 30 thousands SINE elements(Figure 4D).

**Figure 4:**
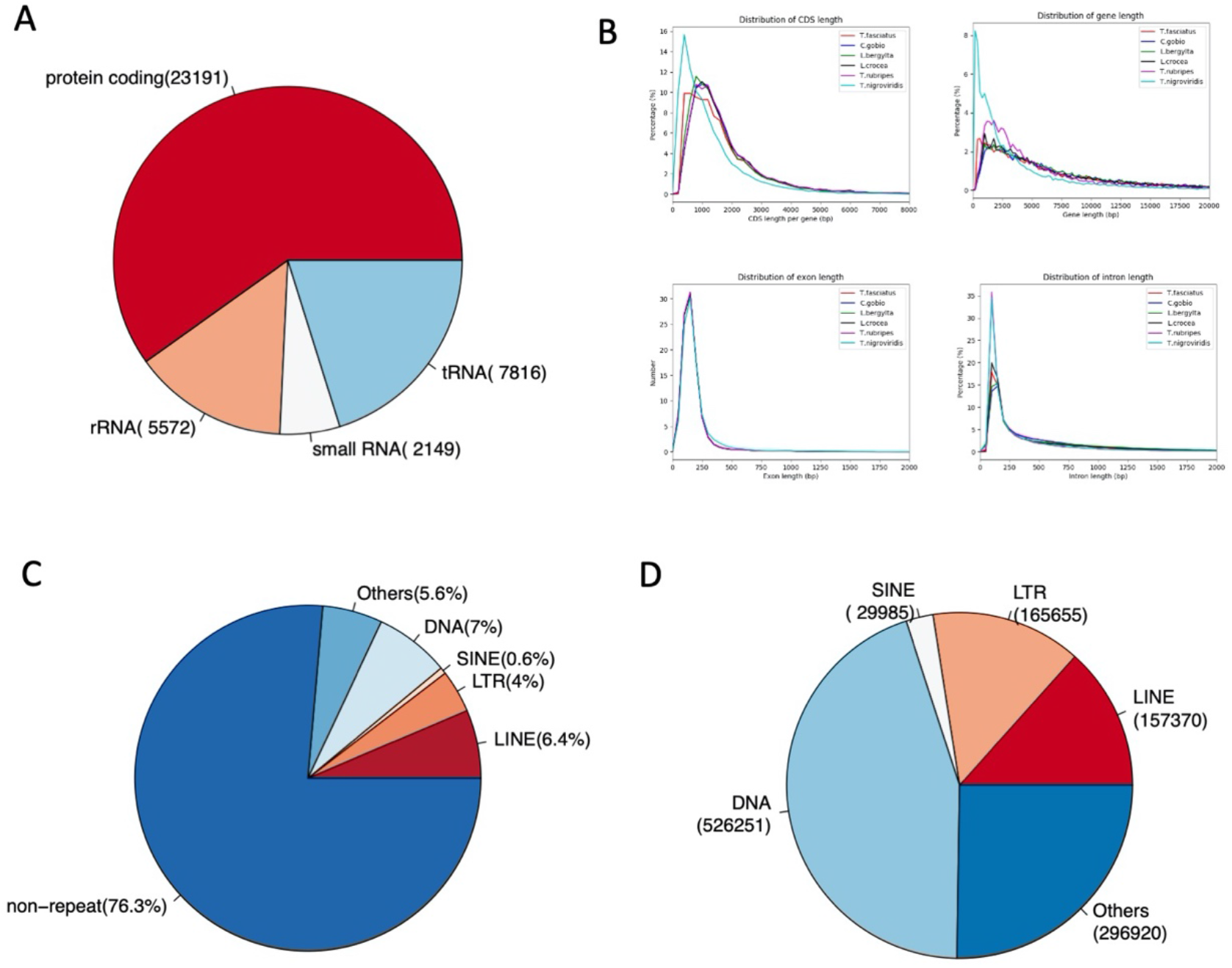
Summary about the genome annotation for *Trachidermus fasciatus*. (A) Number of protein coding and non-protein coding genes. (B) Comparing gene annotation length distributions among close species. (C) Percentage of genomes occupied by repetitive elements. (D)Number of each repetitive element classes.

## Methods

### Tissue extraction and sequencing

One-year old live *Trachidermus fasciatus* fish was collected for tissue extraction. The fish was killed by head smashing, and then quickly immersed on the ice. Eight different organs were dissected for RNA sequencing, including muscle, heart, skin, liver, gill, spleen, gall bladder and stomach. Muscle sample was also used for DNA sequencing (nanopore and Illumina sequencing) and Hi-C sequencing (Illumina sequencing). The tissue samples were stored in liquid nitrogen before sequencing. *Trachidermus fasciatus* genomic DNA was extracted by QIAGEN® Genomic DNA extraction kit (Cat#13323, QIAGEN) following standard protocol suggested by the manufacturer, and RNA sequencing library was constructed by Illumina TruSeq RNA library Preparation Kit. Nanopore sequencing was carried on Nanopore GridION X5/PromethION sequencer, while Illumina sequencing (DNA, RNA, Hi-C) was carried on Illumina HiSeq platform. Finally, Hi-C library preparation was done by following the protocol reported previously[17].

### Genome de novo assembly

NextDenovo was used for genome assembly, including sequencing error correction, preliminary assembly and genome polishing. Basically, NextCorrect module was used for raw reads correction and consensus sequence (CNS) extraction. NextGraph module was used for preliminary assembly, and Nextpolish[18] module was used for genome polishing. At the genome polishing stage, nanopore reads were used repetitively three times and Illumina sequencing reads were used repetitively four times for genome correction. Seed cutoff was set at 38Kbp, and reads cutoff was set at 1Kbp for the NextDenovo genome assembly, and default parameters were used for other settings.

### Genome assembly quality evaluation

Four evaluation metrics were applied for the quality evaluation of the genome assembly, including completeness of the genome, genome accuracy and consensus, GC proportion and sequencing depth distribution (GC-depth analysis), and genome contaminations. Both of BUSCO[19] and CEGMA[20] were used for genome completeness evaluation. BUSCO evaluated the completeness of the assembly by matching it with the ortholog genes from OrthoDB[21] database, and the evaluation of CEGMA was done by comparing the evolutionarily conserved core protein coding genes in eukaryotes (248 core genes). In order to assess the assembled genome sequence accuracy and consensus, Illumina sequencing reads was mapped onto the genome by bwa. Samtools and bcftools[22] were used for single nucleotides polymorphism (SNP) and insertion deletion (Indel) calculation. The percentage of homogeneous SNPs were considered as the single nucleotide error rate of the assembled genome. For the GC-depth analysis, the nanopore sequencing reads were mapped onto the genome assembly by minimap2[23], and the GC content proportion and long reads coverage were calculated for each sliding windows (size of 10kb) of the assembled genome. Finally, the assembled genome was compared to the sequences from Nucleotide Sequence Database (NT, ftp.ncbi.nih.gov/blast/db) to examine the contamination from other species. In detail, the genome was divided into 1Mb bins and then aligned with the NT sequences by blastn[24] software. The mapping statistics was summarized based on the results from each bins.

### Hi-C guided Chromosome level scaffold assembly

The raw paired-end Hi-C reads were preprocessed by fastp[25] for adapter trimming, low quality reads filtering (only keep reads with Phred Score > 15, and number of Ns in the reads less than 5).Each pair of the clean reads were mapped onto the assembled genome by bowtie2[26] (version: 2.3.2, parameters: -end-to-end, --version-sensitive - L 30). For the reads that cannot be mapped onto the genome, DpnII restriction endonuclease recognition sequence pattern GATC was searched, and the reads were cut by restriction sites and used for further mapping. Each pair of the uniquely mapped reads were merged for further analysis. LACHESIS[15] (parameters: CLUSTER MIN RE SITES = 100; CLUSTER MAX LINK DENSITY=2.5; CLUSTER NONINFORMATIVE RATIO = 1.4; ORDER MIN N RES IN TRUNK=60; ORDER MIN N RES IN SHREDS=60;) was used to obtain chromosome-level scaffolds based on the primary assembly and the Hi-C reads mapping information.

### Gene and repetitive element annotation

Based on the RNA sequencing data from eight tissues, the assembled genome and public homolog protein sequences, *Trachidermus fasciatus* genome was annotated at different levels, including the annotations of repetitive elements, non-protein coding RNAs (ncRNA), and protein coding genes. Firstly, RepeatMasker[27] (www.repeatmasker.org) was applied to annotate the repetitive elements, and the repeats masked genome was further used for gene annotation. For the protein coding gene annotation, Evidence Modeler (EVM)[28] was used to integrate the annotation results from three methods, including transcriptome prediction by PASA[29], homolog protein prediction by GeMoMa[30], and de novo gene prediction by AUGUSTUS[31]. Furthermore, Infernal[32] was used to predict ncRNAs, and tRNAscan-SE[33] was chosen to predict tRNAs.

### Data availability and software detail

The genome sequence of Trachidermus fasciatus had been deposited in the Genome Warehouse in National Genomics Data Center [34], Beijing Institute of Genomics (BIG), Chinese Academy of Sciences, under accession number GWHACFF00000000 that is publicly accessible at https://bigd.big.ac.cn/gwh.R was used for graphical plotting, and the source and version of the software used is listed as below.

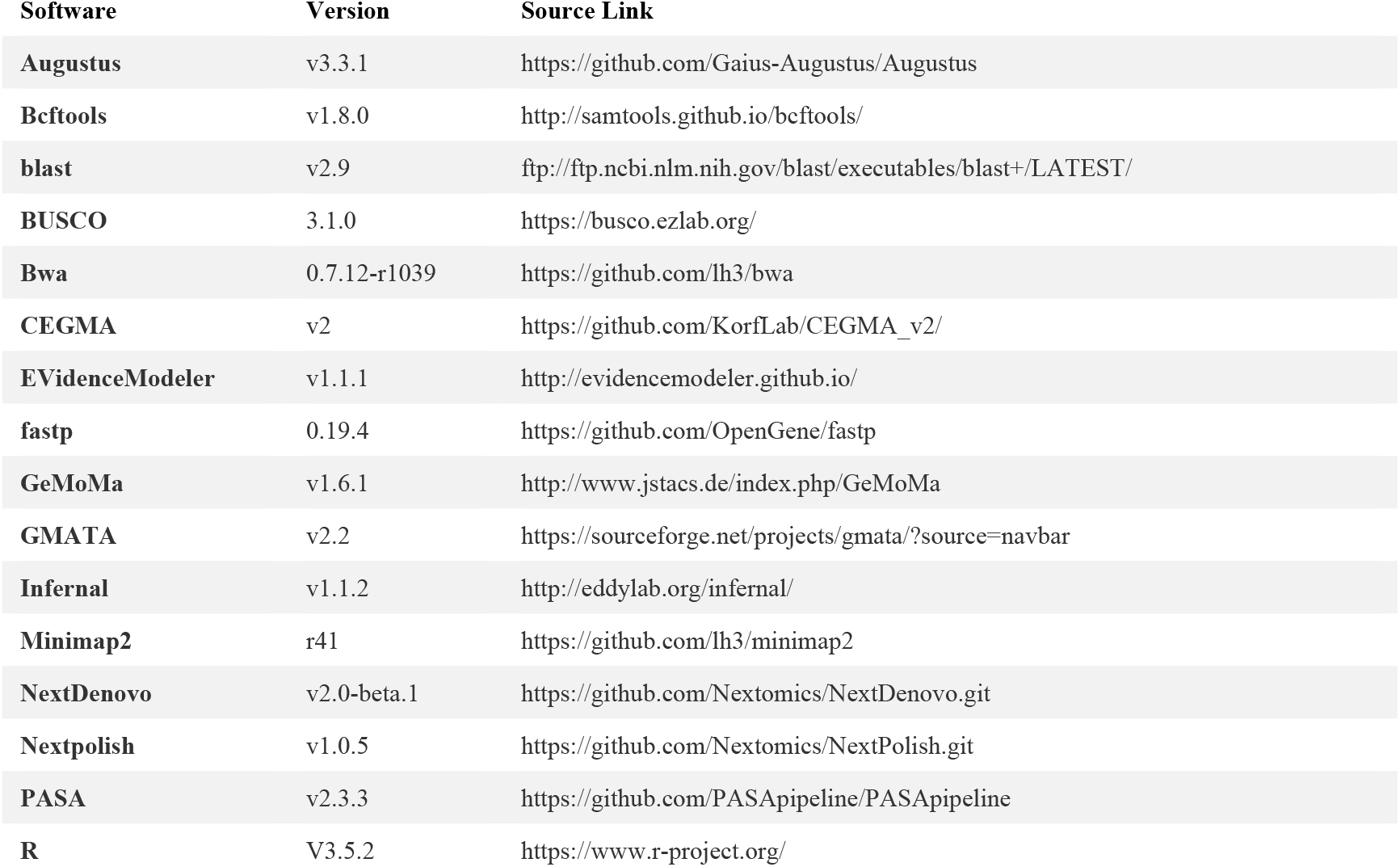
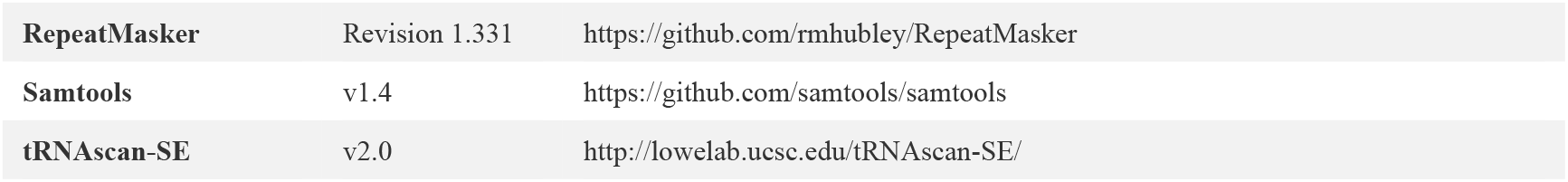

## Discussion

In this study, we provided the first complete genome assembly for roughskin sculpin fish *Trachidermus fasciatus*. Due to overfishing and environmental destruction, the population of roughskin sculpin fish is currently under treat in China, and also had long been listed as Class II protected animal. Our study might be important for future studies for the protection and domesticated culturing of *Trachidermus fasciatus*. Besides providing the first genome reference, we also predicted gene structures for this species, and in total annotated 25,741 genes. Genome annotation information is important for future genetics study of this species, which provided a rich genetic resource for phenotypical and ecological studies. Although we had made the genome assembly and gene annotation publicly available for the research community, the searching and visualization of the genome and gene information for *Trachidermus fasciatus* is still lacking. In the future, we will develop software and databases to improve the accessibility of the genome-wide information for this species.

## Acknowledgements

This study was supported by the grants from National Natural Science Foundation of China (31900484 to Gangcai Xie; 81870359, 2018YFA0801004 to Dong Liu), Natural Science Foundation of Jiangsu Province (BK20190924 to Gangcai Xie; BK20180048, 17KJA180008 and BRA2019278 to Dong Liu), start-up fund for doctoral research of Nantong Science and Technology College (NTKY-Dr2017001 to Feng Lv), Jiangsu Qing and Lan project(2018), Nantong Municipal Science and technology program (JCZ18012 to Feng Lv) and the Open Program of Key Laboratory of Cultivation and High-value Utilization of Marine Organisms in Fujian Province(2019 fjsccq08 to Feng Lv).

## Author Contributions

GX, DL, and JW supervised and designed this project. GX, DL, JW and HH wrote the manuscript. GX, MS analyzed the data. XZ and FL did the experiments.

